# m^6^A RNA methylation regulates promoter proximal pausing of RNA Polymerase II

**DOI:** 10.1101/2020.03.05.978163

**Authors:** Junaid Akhtar, Yoan Renaud, Steffen Albrecht, Yad Ghavi-Helm, Jean-Yves Roignant, Marion Silies, Guillaume Junion

## Abstract

RNA Polymerase II (RNAP II) pausing is essential to precisely control gene expression and is critical for development of metazoans. Here, we show that the m^6^A RNA modification regulates promoter-proximal RNAP II pausing. The m^6^A methyltransferase complex (MTC), with the nuclear reader Ythdc1, are recruited to gene promoters. Depleting the m^6^A MTC leads to a decrease in RNAP II pause release and in Ser2P occupancy on the gene body, and affects nascent RNA transcription. Tethering Mettl3 to a heterologous gene promoter is sufficient to increase RNAP II pause release, an effect that relies on its m^6^A catalytic domain. Collectively, our data reveal an important link between RNAP II pausing and the m^6^A RNA modification, thus adding another layer to m^6^A-mediated gene regulation.

## Introduction

N6-methyladenosine (m^6^A) has recently emerged as the most prevalent mRNA modification in eukaryotes, controlling various developmental, cellular, and molecular processes such as pre-mRNA splicing, alternative polyadenylation, mRNA decay, and translation ^1–3^. Genome-wide mapping of m^6^A in vertebrates revealed a preferred enrichment at specific subsets of sites centered mostly around stop codons, last exons and 3’-untranslated regions (UTRs) of many mRNAs as well as, to a smaller extent, at 5’ UTRs ^4,5^. m^6^A is deposited on mRNA by a multi subunit complex including methyltransferase-like 3 (METTL3), methyltransferase-like 14 (METTL14) ^6^, and Wilms tumor 1-associating protein (WTAP in vertebrates and Fl(2)d in *Drosophila*). This complex is commonly referred to as the m^6^A “writer”. The downstream function of m^6^A is mediated by YTH domain containing protein families, known as “readers”. Ythdc1 is currently the only known nuclear reader protein in *Drosophila*. The majority of m^6^A is deposited co-transcriptionally ^7^, nevertheless the function of this modification in the regulation of RNAP II transcription remains to be identified.

## Results

To address whether m^6^A can directly regulate transcription, we first precisely mapped m^6^A MTC recruitment to chromatin. We performed ChIP-seq experiments with HA-tagged core components of the m^6^A writer complex (Mettl3, Mettl14, and Fl(2)d) and of the m^6^A nuclear reader Ythdc1 in *Drosophila* S2R+ cells. We observed a significant enrichment of m^6^A complex components on chromatin, particularly at promoters (Fig. 1a-c Fig. S1 a-f). m^6^A complex components are similarly enriched at the promoter of intronless genes (Fig. 1b), suggesting that recruitment to chromatin is independent of the reported role of m^6^A in splicing. This enrichment at promoters is consistent with a considerably higher number of m^6^A sites detected at 5’UTRs in *Drosophila* (Fig. S1 g, and ^8^). Importantly, there is a high degree of overlap between the different m^6^A complex components binding sites (Fig. 1d, Fig. S1 h), and the proportion of promoter-bound regions is higher at co-bound sites (Fig. 1e and Fig. S1 c-f). These co-bound regions are enriched for genes associated with developmental and metabolic processes and with cellular response to stimulus (Fig. S1 i). Besides, genes co-bound by the m^6^A MTC tend to be more highly expressed than unbound genes (Fig. 1f), suggesting that m^6^A is preferentially deposited on highly expressed genes. Interestingly, recruitment of the m^6^A MTC appears to be RNA-dependent, as RNAseT1 treatment results in reduced binding (Fig. 1a-c), and blocking transcription initiation with triptolide causes the loss of Mettl3 binding at its target genes (Fig. 1g). However, blocking elongation with 5,6-dichlorobenzimidazole 1-β-d-ribofuranoside (DRB) does not affect Mettl3 binding (Fig. 1g). This RNA-dependent recruitment also requires Mettl3 catalytic function. Indeed, mutating the DPPW motif of Mettl3, essential for its enzymatic activity ^9^, abolished Mettl3 recruitment to chromatin (Fig. 1h). Together, these results strongly suggest that components of the m^6^A MTC and nuclear reader are predominantly recruited to promoters of highly expressed genes in a transcription dependent manner. Moreover, this recruitment requires transcriptional initiation but takes place prior to elongation.

**Fig. 1.**
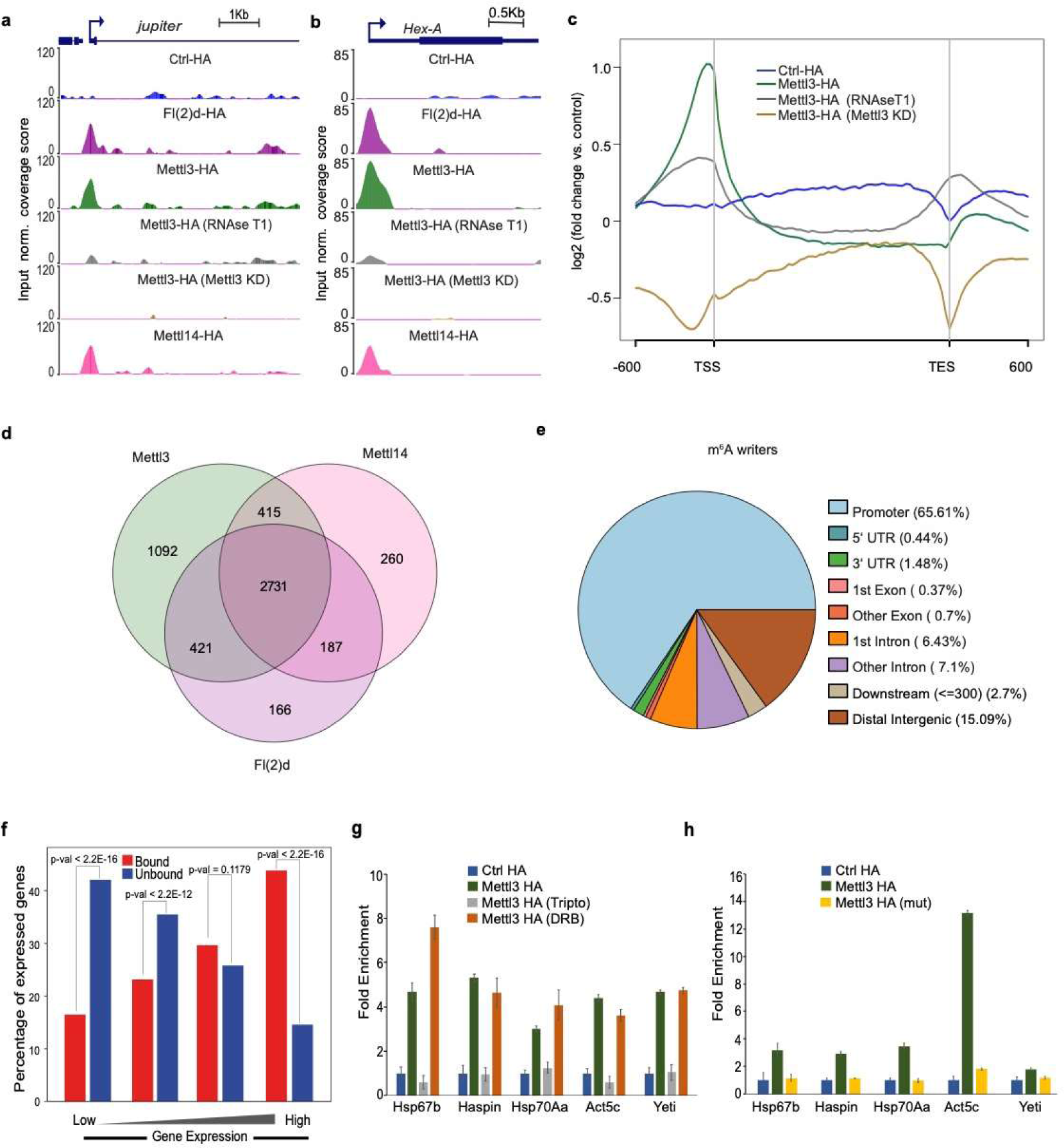
The m^**6**^A complex (MTC) gets recruited to chromatin in an RNA dependent and splicing-independent fashion. **(a, b)** ChIP-Seq tracks from S2R+ cells transfected with HA-tagged MTC components, along with Mettl3 knockdown and RNAse treatment, and control. Shown here are recruitments of MTC to **(a)** an intron-containing and **(b)** intronless gene. **(c)** Averaged metagene profiles of ChIP-Seq performed with HA-tagged Mettl3 and indicated controls with standard error of mean (SEM). **(d)** Venn diagram showing the overlap between genes bound by all the components of the m^6^A MTC. **(e)** Pie chart depicting the distribution of annotated genomic features of all m^6^A MTC components **(f)** Histogram showing percentage of genes bound by m^6^A MTC amongst different quartiles of expressed genes, from least to most expressed. See method section for details. **(g)** ChIP-qPCR analysis of Mettl3-HA occupancies at the promoter of indicated genes in control and after treating with triptolide (initiation inhibitor) and DRB (elongation inhibitor). Error bars indicate the standard deviation from the mean in three biological replicates. **(h)** ChIP-qPCR analysis of HA-tagged Mettl3 and catalytically-dead Mettl3 (DPPW mutant) occupancies at the promoter of indicated genes. Error bars indicate the standard deviation from the mean in three biological replicates.

To further explore the potential effect of m^6^A on transcription, we performed RNAP II ChIP-seq assays before and after individually depleting m^6^A MTC components in S2R+ cells. The knockdown (KD) of m^6^A MTC components resulted in increased RNAP II occupancy at promoters, concomitant with a decrease along the gene body (Fig. 2a-c, Fig. S2 a-c). The deposition of m^6^A MTC also follows the pattern of Pol II occupancy (Fig. 2b). To investigate the effect on RNAP II distribution in greater detail, we calculated the ratio of RNAP II between the gene body and the promoter, hereafter referred as the “RNAP II release ratio” (PRR) (Fig. S2 d). Knocking down individual m^6^A writer components results in a significant decrease of the PRR distribution (Fig. 2d, (Kolmogorov-Smirnov test of distributions, with p-values smaller than 2.2 x 10^−14^ for all the KD conditions)). Importantly, genes displaying the lowest PRR are significantly enriched for m^6^A MTC binding (Fig. 2e), suggesting a preferential recruitment of m^6^A MTC on paused genes. Paused genes (i.e. genes with a low PRR) also tend to be more affected by the KD of one of the m^6^A complex components (here Mettl3) at promoters that are co-bound by all m^6^A MTC components compared to unbound promoters, consistent with a direct role of Mettl3 on transcription (Fig. 2f). This observation was also true for other members of the m^6^A MTC (Fig. S2 e-h), as well as for the nuclear reader protein Ythdc1, indicating that the effect is mediated through m^6^A RNA modification (Fig. S2 b-c and S2 h). In conclusion, our results provide the first evidence of a direct positive impact of the m^6^A RNA modification on RNAP II pause release.

**Fig. 2.**
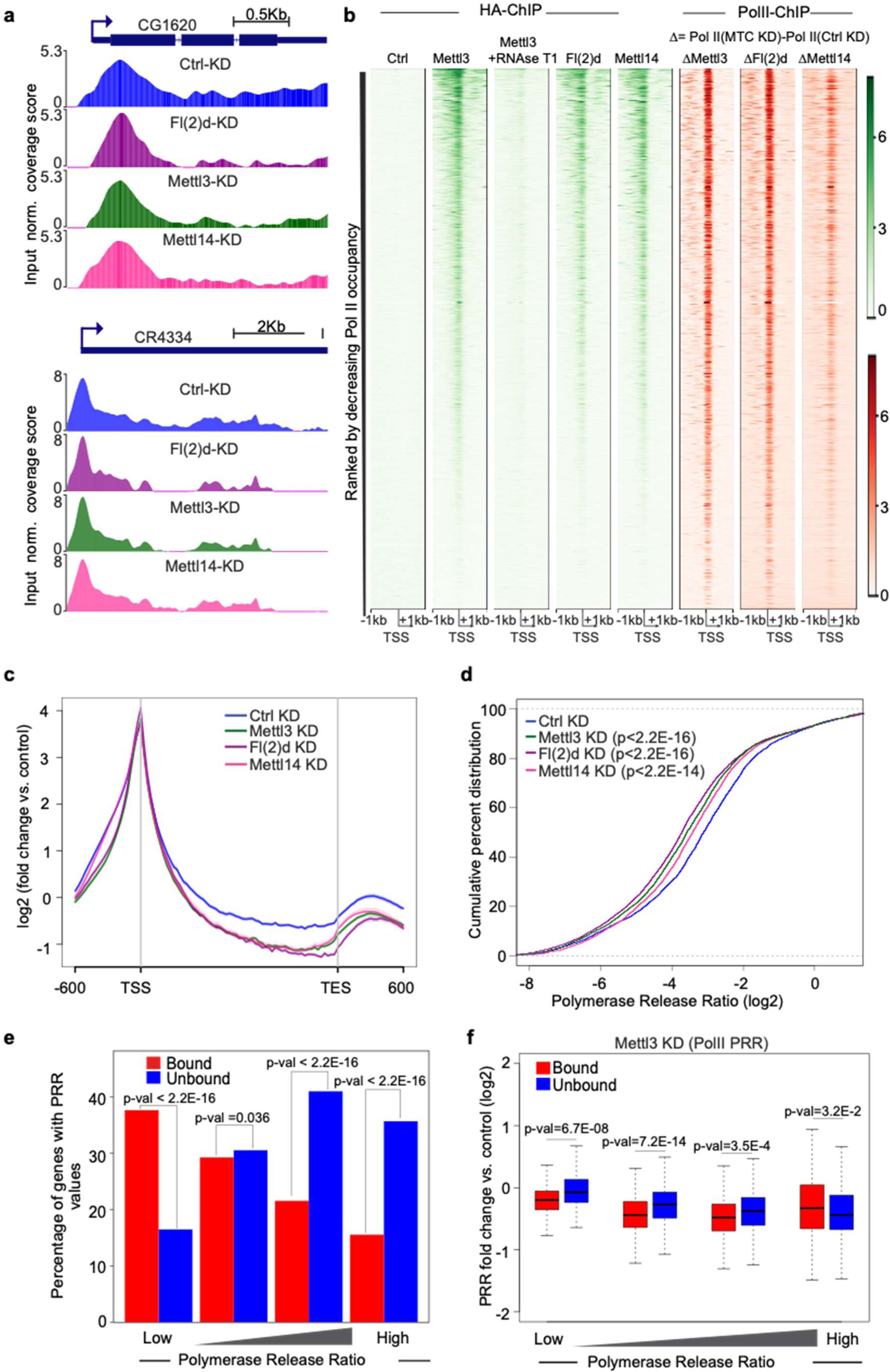
Binding of the m^**6**^A complex to promoters modulate the release of RNAP II from paused state. **(a**) Track examples of total Pol II ChIP-Seq after control or m^6^A MTC knockdowns, as indicated in the labels. The tracks are averaged after input and “spike-in” normalization. **(b)** Heatmaps of HA-tagged m^6^A MTC and change in RNAP II upon m^6^A complex KD, centered at the TSS (−1kb to +1kb), and sorted by decreasing RNAP II occupancy. The color labels to the right indicate the levels of enrichment. **(c)** Metagene profiles of averaged RNAP II occupancies after “spike-in” normalization in control and m^6^A MTC depleted cells with SEM for all the RNAP II bound genes. Log2 fold changes against input are shown on Y-axis. **(d)** The empirical cumulative distribution function (ECDF) plot of computed PRR in control and m^6^A MTC knockdown conditions, after “spike-in” normalization. **(e)** Histogram showing percentage of m^6^A MTC-bound genes amongst different PRR quartiles in control condition, sorted according to the lowest PRR (most paused) to the highest PRR (least paused) level. See method section for details. **(f)** Boxplot showing change in PRR upon Mettl3 knockdown in different PRR quartiles as in **(f)** and further separated based on binding of MTC.

To characterize the role of m^6^A in RNAP II pause release, we investigated the influence of the m^6^A MTC on RNAP II Ser2 phosphorylation (Ser2P), a hallmark of transcription elongation. We performed ChIP-seq experiments with RNAP II Ser2P-specific antibodies in m^6^A MTC depleted S2R+ cells. The most striking effect of m^6^A MTC depletion is a significant decrease in Ser2P level along the gene body, indicating that the transition from paused state to productive elongation is affected (Fig. 3a-c, and Fig. S3 a). We further validated this result using the λN-BoxB system, which mimics the RNA-dependent binding of Mettl3 at a heterologous gene locus. We tested the transcriptional effect of ectopically tethering Mettl3 to this construct ^10^ and showed that the ectopic binding of Mettl3 alone at the luciferase reporter is sufficient to decrease RNAP II and Ser2P enrichment at the promoter and increases it along the gene body (Fig. 3d). As this effect is observed using the Luciferase CDS, we assume that it does not rely on splicing. Next, we tested whether the transcriptional regulation by Mettl3 was dependent on its enzymatic activity on the RNA. Indeed, in contrast to wild type Mettl3, tethering the catalytically dead Mettl3 ^9^ using λN-BoxB heterologous system does not affect the transcription of the luciferase reporter (Fig. 3e). Therefore, the RNA catalytic activity of Mettl3 is essential for its role in transcriptional regulation. To further validate the effect of m^6^A on RNAP II pause release, we sequenced transiently synthesized nascent RNAs using DRB-4SU-Seq, an approach particularly suited to study RNA elongation rates and initiation frequencies through the reversible inhibition of transcription by DRB and the tagging of nascent transcripts with 4-thiouridine (4SU) (Fig. S3 b, and method section). To mitigate any confounding factors, we restricted our analysis to genes longer than 15kb and reads at least partially mapping to introns ^11^. The depletion of Mettl3 or Ythdc1 results in decreased level of nascent RNAs 2 minutes after DRB removal (Fig. 3f and 3g). This decrease in nascent transcripts and Ser2P level along the gene body upon depletion of m^6^A complex components suggests that the transition of RNAP II from a paused state into productive elongation is mediated through m^6^A deposition on mRNAs. Together, these results strongly support a role of the m^6^A MTC in directly regulating the release of RNAP II from a paused state in an m^6^A RNA dependent fashion.

**Fig. 3.**
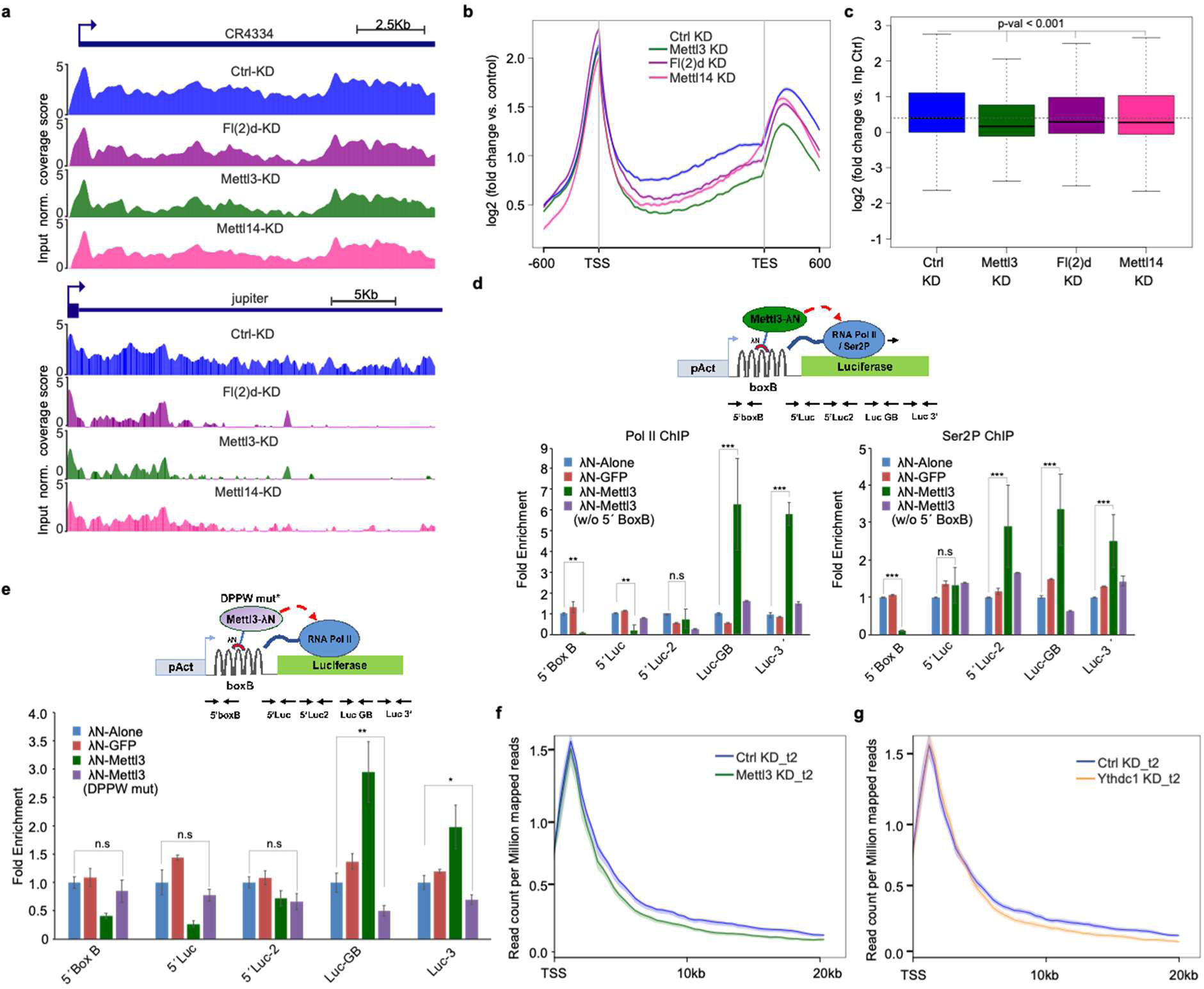
Loss of m^**6**^A MTC leads to decrease in transcriptional elongation. **(a**) Track examples of Ser2P after control or m^6^A MTC knockdowns. The tracks are averaged after input and “spike-in” normalization. **(b)** Metagene profiles of averaged Ser2P occupancies after “spike-in” and input normalization in indicated KDs. Log2 fold changes against input are on Y-axis. **(c)** Boxplot showing Ser2P occupancy on the gene bodies in indicated KDs after input normalization. **(d, e)** (Top) Scheme: BoxB-λN tethering assay. BoxB sequences (hairpins) were inserted upstream of Firefly luciferase (green). The λN peptide (red) was fused to Mettl3 (green), GFP or alone, and transfected into S2R+ cells with the modified luciferase as well as Renilla construct. (Bottom) Quantification of the ChIP enrichment. The enrichment of RNAP II (left) and Ser2P (right) at the indicated loci after normalizing against a negative locus and Renilla **(d). (e)** Similar experimental set up as in **(d)**, only that a catalytically-dead Mettl3 was used and ChIP was performed for RNAP II. The enrichment for three independent biological replicates is shown. **(f, g)** Metagene profile of nascent RNA from DRB treated 4sU-Seq data in control and knockdown conditions with standard error of mean for all the expressed genes longer than 15 kb, based on the average of two independent biological replicates. Shown are the profiles after 2 minutes of DRB removal upon Mettl3 **(f)** and Ythdc1 **(g)** knockdowns. Averaged read counts per million of mapped reads of two independent biological replicates from 4sU-Seq are shown on Y-axis while X-axis depicts genomic coordinates.

Recent evidence in mammals suggests that H3K36me3 enrichment on chromatin guides m^6^A deposition on mRNAs ^12^. Surprisingly, we did not observe any correlation between the binding of m^6^A components and H3K36me3 enrichment in *Drosophila* (Fig. S3 c-d). Indeed, knocking down SetD2, the histone methyltransferase responsible for lysine 36 methylation of histone H3, only caused a moderate reduction of m^6^A complex binding at particular loci, confirming the existence of a H3K36me3-independent recruitment of the m^6^A MTC on promoters (Fig. 4 d). Instead, *de novo* motif analysis at m^6^A MTC co-bound peaks revealed DREF (p < 1e-178), GAF (p < 1e-169) and M1BP (p < 1e-250) as the most enriched motifs, altogether represented in ∼ 40% of the MTC peaks (Fig. S4 a). These factors were previously shown to regulate RNAP II pausing in independent studies ^13,14^, reinforcing the connection between m^6^A and RNAP II pausing. We further investigated the correlation between the deposition of GAF and M1BP and components of the m^6^A writer complex using ChIP-seq datasets from S2R+ cells ^13,14^. Genome browser tracks and cumulative metagene analysis of GAF and M1BP profiles revealed correlated distribution patterns of these factors with m^6^A MTC (Fig. S4 b-c). Similarly, there is a considerable overlap between genes co-bound by the m^6^A complex components and genes bound by either GAF or M1BP (Fig. S4 d). These data suggest that the deposition of the m^6^A RNA modification at promoter in *Drosophila* is not guided by the H3K36me3 chromatin modification, but instead depends on other features that remain to be uncovered.

**Fig. 4:**
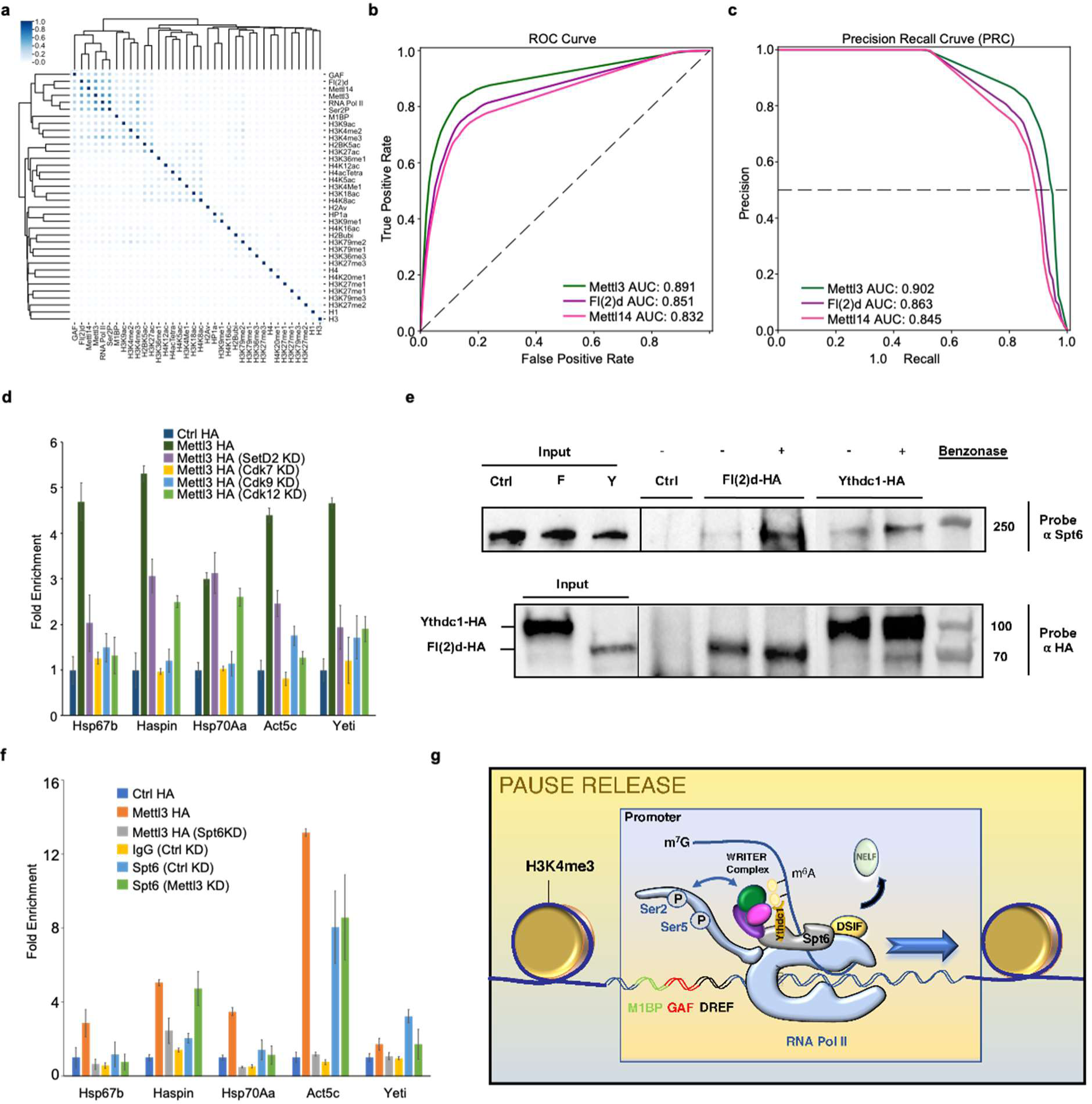
Mettl3 promoter recruitment depends on Spt6, and CTD phosphorylation. **(a)** Clusterogram of m6A complex datasets together with the publicly available datasets of indicated histone marks, M1BP and GAF. The clusterogram represents an agglomerative clustering with average linkage based on similarity indices calculated on the overlapping genomic windows of 1kb, derived from the signal in the peak file. The legend in the left indicates the distance based on Jaccard Index. **(b, c)** ROC **(b)** and PRC **(c)** curves describing the performance of Random Forest classification models in predicting the recruitment of HA-tagged m^6^A MTC components; used to define the class values. The RNAP II and Ser2P datasets from this study and published datasets of GAF and M1BP were used by the model as classification features together with the histone modification datasets in S2R+ cells from modENCODE consortium. **(d)** ChIP-qPCR analysis of Mettl3-HA occupancies at the promoter of indicated genes, in labeled knockdown conditions. Error bars are from three biological replicates. **(e)** Co-immunoprecipitation using anti-Spt6 antibody from cell extracts expressing either HA-Fl(2)d, HA-Ythdc1 or HA alone (Ctrl), revealed with Spt6 and HA antibodies. Note that Fl(2)d and Ythdc1 interacts with Spt6 without a DNA or RNA intermediate, as treating the lysate with benzonase does not result in the loss of the interaction. **(f)** ChIP-qPCR experiments showing recruitment of Mettl3 at the promoter of indicated genes, in control and Spt6 knockdown conditions. Note that Mettl3 recruitment was dependent on Spt6 as knocking down Spt6 resulted in strong decrease in Mettl3 recruitment. However, Spt6 recruitment was resistant to Mettl3 KD. Error bars indicate the standard deviation from the mean of three biological replicates. **(g)** Model: The regulation discovered here suggests a mechanism where transcription dependent recruitment of the m^6^A complex components positively feedbacks on the transcription machinery to promote release of RNAP II from paused state, through its interaction with RNAP II and Spt6.

To further identify the features predictive of m^6^A MTC binding, we gathered publicly available datasets for various histone marks in S2R+ cells. We combined these datasets with the previously described m^6^A MTC, RNAP II, Ser2P, GAF and M1BP ^13,14^ datasets to generate a similarity Jaccard index calculated on sets of genomic windows. The heatmap shows that GAF, M1BP, RNAP II and Ser2P binding sites cluster with the m^6^A MTC bound regions (Fig. 4 a). We then used the same datasets as features in a machine learning approach using the Random Forest classification algorithm to train models that predict the binding of m^6^A MTC. The receiver operating characteristics (ROC) curves show an 80% true positive rate (TPR) with lower (< 20%) false positive rate (FPR) and high Area Under the Curve (AUC > 0.83) (Fig. 4b). Furthermore, the precision recall curves (PRCs) confirm high predictive performance in feature-dependent recalling of m^6^A complex component binding (Fig. 4c). Interestingly, among the top 10 features used by the Random Forest models, RNAP II, Ser2P and binding of pause factors at promoters are the most important, followed by activating histone marks (Fig. S4 e-g). Consistent with this, ChIP-qPCR experiments upon Cdk7 ^15^, Cdk9 ^16^, or Cdk12 KD ^17^, that leads to decrease in Ser2P level, also caused a strong decrease of Mettl3 occupancy at interrogated promoters (Fig. 4d). These results validate the machine learning performance and suggest a role of RNAP II phosphorylated forms in recruitment of m^6^A MTC. Collectively, these data suggest that a feedback mechanism exists between m^6^A and the transcription machinery.

To identify the mechanism behind m^6^A-dependent RNAP II pause release, we performed mass-spectrometry analysis after oligo pull-down using a m^6^A methylated RNA bait containing a strong consensus motif. We compared the interactome of this oligo with a non-methylated one and identified Spt6 among the top m^6^A RNA interactors (Table 1). Spt6 is a histone chaperone protein associated with the elongation complex that is co-localized with different forms of RNAP II in *Drosophila* and is implicated in transcriptional regulation ^18^. Co-immunoprecipitation experiments performed in the presence of a nuclease confirmed the interaction between Spt6 and the m^6^A complex components Fl(2)d and Ythdc1, without any nucleic acid intermediate (Fig. 4e). We next asked whether the recruitment of the m^6^A MTC was dependent on Spt6. ChIP-qPCR analysis on m^6^A MTC bound loci shows a significant loss of Mettl3 promoter occupancy upon Spt6 KD (Fig. 4f). On the contrary, Spt6 binding on chromatin is not affected by Mettl3 depletion (Fig. 4f), indicating that Spt6 co-occupies Mettl3-bound promoters in a Mettl3-independent manner. Together, these data provide evidence that m^6^A MTC recruitment to the chromatin at least partially depends on Spt6 binding. This is also consistent with its observed effect on RNAP II pause regulation, and the transition to elongation.

Promoter-proximal RNAP II pausing is a widespread phenomenon in metazoans, and is a key step in the regulation of gene expression. Our data uncovered a mechanism where transcription-dependent recruitment of the m^6^A complex components positively feedbacks on the transcription machinery to promote release of RNAP II from the paused state (Fig. 4g). While we provide the first evidence for a role of m^6^A in RNAP II pausing, this link is not uncalled for. First, m^6^A complex components and pause factors are both found at gene promoters, and the most prevalent paused genes (i.e. developmentally-regulated and stress-associated genes) tend to also be bound by the m^6^A MTC at their promoter. Second, the m^6^A RNA modification plays a critical role during cell fate decision in early mouse development and in stem cell differentiation ^19–21^. Interestingly, these same processes are highly subjected to RNAP II pausing and might involve m^6^A RNA for their precise control. Finally, m^6^A was recently shown to indirectly inhibit transcription by regulating the stability of chromatin-associated RNAs ^22^. While this result might seem to contradict our findings (which rather point towards a positive action of m^6^A on transcription regulation), this potential multifaceted and context-dependent regulation is reminiscent of a similar action of the Polymerase associated factor 1 (PAF1). Indeed, depending on the cell type, PAF1 is required either for stabilizing RNAP II pausing or for promoting its pause release ^23,24^. The m^6^A-mediated control of RNAP II pausing adds a novel layer to an already highly regulated process. Such regulation is akin to the well-established transcriptional checkpoint associated with the capping of nascent RNA and splicing ^25^.

## Acknowledgments

We thank the EMBL Genomics Core Facility (Heidelberg, Germany), especially Vladimir Benes, for all the sequencing runs. We thank Nastasja Kreim (IMB, Germany) for the constant support and discussions. We also thank Sebastian Molina Obando (Silies lab), Dr Philippe Arnaud (GReD institute) and Dr Guillaume Lavergne (Jagla’s lab) for comments on the manuscript.

## Author contributions

J.A. conceived the idea and started the study. J.A and G.J designed and performed the experiments. G.J and J.A. supervised Y.R. and S.A. in genome-wide datasets analysis. G.J. with Y.G.H. performed the LC–MS/MS quantification of m^6^A proteome. J.Y.R. and M.S. gave inputs, J.A. and G.J. wrote the manuscript with inputs from all authors.

