## Supplementary material for "m^6^A RNA methylation regulates promoter proximal pausing of RNA Polymerase II": Supplemntary Materials (from another version of this preprint: version 1, 2020-03-05)

### Materials and Methods

#### Cloning

The plasmids used for chromatin immunoprecipitation (ChIP) and co-IP assays in *Drosophila* S2R+ cells were constructed by cloning the corresponding cDNA in the pPAC vector with N-terminal Flag – 3X HA tag, using EcoRV and NotI sites. The lambdaN and Box-B plasmids are derived from earlier studies. The lambdaN plasmids were created by replacing the Mago CDS with Mettl3 CDS and Mettl3 mutant CDS using EcoRV and NotI sites. For Mettl3 mutant CDS, the DPPW motif was mutated to GAAG.

#### RNA Isolation and RT-PCR

RNA was extracted from cells using Trizol reagent, following the manufacturer's protocol. For reverse transcription, cDNA was synthesized using MMLV reverse transcriptase (Promega, Cat No-M1701). Relative abundance of transcripts relative to Rpl15 was calculated by the $2^{\Delta}$ Ct method. PCR primers used for semi-quantitative and real time PCR are listed in Table S2-S4.

#### Cell Culture, RNAi and Transfection

*Drosophila* S2R+ cells were cultured in Schneider Cell's Medium (PAN Biotech, Cat No- P04-91500) supplemented with 10% FBS (Sigma, Cat No- 12103C) and 2% Penicillin/Streptomycin (GIBCO, Cat No- 15140122). The plasmids expressing various transgenes were transfected with Effectene transfection reagent (Qiagen, Cat No-301425), following manufacturer's protocol. For knockdown experiments, dsRNA was synthesized overnight at 37 °C using Hi-Scribe T7 transcription kit (NEB, Cat No-E2040). dsRNA was transfected in S2R+ cells by serum starvation for 6 hours. The treatment was repeated three times and cells were harvested 5 days after the first treatment. For Cdk7, Cdk9, Spt6 and Cdk12 knockdowns, the treatment was repeated twice, and cells were harvested 5 days after the first treatment. All the dsRNA knockdown efficiencies have been validated by RT-qPCR (Fig. S5). The primers used for generating dsRNA are listed on Table S1. Cells were treated with 5,6-dichlorobenzimidazole 1-β-d-ribofuranoside (DRB) from Sigma (D1916) at 300 μM, dissolved in water, for 5 h to block elongation. The triptolide (Sigma, Cat No- T3652) treatment was performed for 6 h at 10 μM.

**Antibodies**

The following antibodies were used in this study. For total Pol II ChIP, RBP1 (Diagenode Cat No-15200004) was used. Ser2P ChIP was performed using ab5095 (Abcam) and for H3K36me3 ab9050 (Abcam) was used. For co-immunoprecipitation experiments, anti-HA (Sigma, Cat No. 11867423001), anti-Spt6 (a gift from John T Lis, Cornell University, Ithaca, NY, USA) and Dynabeads Protein-G (ThermoFisher, Cat No-10003D) were used. For HA-tagged ChIP experiments anti-HA antibody from Santa Cruz was used (Cat. No. sc-7392 X).

**Co-IP Assay and Western Blot Analysis**

For co-IP assay in S2R+ cells, cells were plated in 10 cm cell culture dish, and respective transgenes were transfected using Effectene transfection reagent, according to manufacturer's protocol. 48h post transfection cells were collected, washed once with PBS and re-suspended in swelling buffer (10mM Tris pH 7.5, 2 mM MgCl2, 3mM CaCl2, and protease inhibitors). After incubating 10 min on ice, the suspension was spun at 600g for 10 min at 4°C. After discarding the supernatant the pellet was resuspended in lysis buffer (10mM Tris pH 7.5, 2 mM MgCl2, 3mM CaCl2, 0.5% NP-40, 10% glycerol and protease inhibitors) and centrifuged for 5 min at 600g. Nuclei were resuspended in lysis buffer 2 (40mM HEPES pH7.4, 120mM NaCl, 10 mM MgCl2, 0.4% NP-40 and protease inhibitors) and sonicated in the bioruptor pico (Diagenode) for 5 cycles with 30 Sec "ON/OFF". Protein concentrations were determined using Bradford reagent (BioRad, Cat No-5000006). For IP, 2 mg of proteins were incubated with respective antibody in lysis buffer 2 and rotated head-over-tail O/N at 4 °C. The beads were washed 3x for 10 min with wash buffer ( 40mM HEPES pH7.4, 140mM NaCl, 10 mM MgCl2, 0.4% NP-40, 5% Glycerol and protease inhibitors) and IP proteins were eluted by incubation in 1x Laemmli sample buffer

buffer (Biorad, Cat No-161-0747), 50mM DTT at 95 °C for 10 min. Immunoprecipitated and input proteins were analyzed by western blot, after separating them on 4-15% gradient SDS-PAGE gel (BioRad, Cat No-4561083) and transferred to Nitrocellulose membrane (Biorad, Cat No-1704271). After blocking with 5 % milk in PBT (0,1 % Tween) for 2H at RT, the membrane was incubated with respective primary antibody in blocking solution O/N at 4 °C. Membrane was washed 4 x in PBT for 15 min and incubated 2h at RT with secondary antibody-HRP in blocking solution. Blots were developed using ECL prime (Amersham, RPN2232SK) and visualized using BioRad Chemidoc system.

**ChIP-qPCR and ChIP-Seq**

S2R+ cells were fixed with 1% formaldehyde for 10 min at room temperature and harvested in SDS buffer. The nuclei, collected after centrifuging at 600g for 6 min, were resuspended in RIPA buffer (140 mM NaCl, 10 mM Tris-HCl [pH 8.0], 1 mM EDTA, 1% Triton X-100, 0.1% SDS, 0.1% DOC), and lysed by sonication. The lysate was cleared by centrifugation at 20,000g, and incubated with respective antibodies overnight at 4°C. Antibody complexes bound to protein G beads were washed once with 140 mM RIPA, four times with 250 mM RIPA, once with LiCl buffer and twice with TE buffer for 10 min each at 4°C. DNA was recovered after reverse crosslinking and phenol chloroform extraction, using phase lock tubes and following manufacturer's instructions. After precipitating and pelleting, DNA was dissolved in 30 µl of TE pH 8.0. Control immunoprecipitations were done in parallel with either tag alone or knockdown controls and processed identically. 5 µl of immunoprecipitated DNA was used for checking enrichment with various primer pairs (listed in supplementary table 1) on Applied Biosystem ViiA™ 7 real time machine using SYBR green reagent (Life technologies, Cat No-4367659). To examine whether the changes in Pol II and Ser2P distribution were widespread, we performed ChIP-Seq experiments in control and $m^6A$ complex component's knockdown conditions. To exclude the possibility of changes in Pol II occupancy driven by differences in immunoprecipitation efficiency and technical variance during library preparation in different knockdown conditions, we used yeast chromatin as "spike-in" control (*26*). After validating enrichment, the recovered DNA was converted into libraries using NebNext Ultra DNA library preparation kit, following manufacturer's protocol. DNA libraries were multiplexed, pooled and sequenced on Illumina HiSeq 2000 platform.

**DRB-4sU-Seq**

S2R+ cells were grown in Schneider's Cell Medium with 10% bovine serum supplemented with antibiotics and maintained at 25°C. 5,6-dichlorobenzimidazole 1-β-d-ribofuranoside (DRB) from Sigma (D1916) was used at a final concentration of 300 μM, dissolved in water, for 5 h. 4-thiouridine (4sU) was purchased from Sigma (Cat No-T4509) and used at a final concentration of 100 μM. Control and m6A complex KD was performed as described before. All the samples were labeled for 6 min with 4-thiouridine, and transcription was allowed to proceed after DRB removal for 0, 2, and 10 min along with one non-DRB treated control.

A total of 100 to 130 μg RNA was used for the biotinylation reaction. 4sU-labeled RNA was biotinylated with EZ-Link Biotin-HPDP (Thermo Scientific, Cat No-21341), dissolved in dimethylformamide (DMF, Sigma Cat No-D4551) at a concentration of 1 mg/mL. Biotinylation was done in labeling buffer (10 mM Tris pH 7.4, 1 mM EDTA) and 0.2 mg/mL Biotin-HPDP for 2 h with rotation at room temperature. Two rounds of chloroform extractions removed unbound Biotin-HPDP. RNA was precipitated at 20,000 g for 20 min at 4°C with a 1:10 volume of 5M NaCl and an equal volume of isopropanol. The pellet was washed with 75% ethanol and precipitated again at 20,000 g for 10 min at 4°C. The pellet was left to dry, followed by resuspension in 100 μL RNase-free water. Biotinylated RNA was captured using Dynabeads MyOne Streptavidin T1 beads (Invitrogen, Cat No-65601). Biotinylated RNA was incubated with 50 μL Dynabeads with rotation for 15 min at 25°C. Beads were magnetically fixed and washed with 3× Dynabeads washing buffer. RNA-4sU was eluted with 100 μL of freshly prepared 100 mM dithiothreitol (DTT), and cleaned on RNeasy MinElute Spin columns (Qiagen, Cat No-74204). For the untreated 4sU-Seq version used for calculating polymerase release ratio (PRR), an identical approach was used with following modifications. During the period when biotinylated RNA was incubated with 50 μL Dynabeads with rotation for 15 min at 25°C, RNAse T1 was added in order to fragment RNA to 100bp. Beads were magnetically fixed and washed with 3× Dynabeads washing buffer, as described before. RNA-4sU was eluted with 100 µL of freshly prepared 100 mM DTT, and cleaned on RNeasy MinElute Spin columns (Qiagen, Cat No-74204). Enriched nascent RNAs were converted to cDNA libraries with *Drosophila* Ovation Kit (Nugen- Cat No-7102-32) with integrated ribosomal depletion workflow. Amplified cDNA libraries were pooled, multiplexed, and sequenced on one lane of Illumina Nextseq at 75bp paired end settings.

**$m^6A$ RNA pull-down**

To identify proteins that recognize and bind to $m^6A$ RNA, an oligo-pull-down strategy was implemented. The S2R + cell lysate at the concentration of 0.4 mg/mL was combined with methylated or unmethylated RNAs. The oligos were biotinylated and subjected to Streptavidin beads (Thermo Scientific, Cat No-11205D) pull-downs. The beads were washed 3 times with a RIP washing buffer (Magna RIP Millipore-Cat No. 17-700), supplemented with RNAse inhibitor, and separating them with magnetic tube holder. In each experiment, 50 picomole of biotinylated RNA were added to the beads and incubated for 30 min at RT with shaking and washed 10 times with 0.5ml of RIP washing buffer. After washing the beads, the lysate was added for overnight at 4 °C with head over tails rotation. The beads were further washed with the RIPA buffer for a total of 5 times. The beads were finally resuspended in 30-35µl of ammonium acetate (CH3COONH4) and the samples were sent for mass spectrometry to the IBCP - LYON Mass Spectrometry Service.

Oligos used in these experiments (Gene Link) were 5’ biotinylated and with sequence UGGGCCUGUCAUCGCUAAAAGGA6CUGCUUUUGGGGCUUG for $m^6A$ modification and UGGGCCUGUCAUCGCUAAAAGGACUGCUUUUGGGGCUUG for control.

**miCLIP**

miCLIP was performed following previously described method [2] using 10 µg of purified mRNA from *Drosophila* S2R+ cells and 5 µg of anti-$m^6A$ antibody (Synaptic Systems, Lot# 202003/2-82).

Immunoprecipitations were performed in quadruplicates and as a control one immunoprecipitation was performed where UV-crosslinking was ommited. All steps of library preparation were performed as previously described with minor modifications [3], and would be published elsewhere. Libraries were sequenced on an Illumina NextSeq500.

**Computational Analysis**

**ChIP-Seq:** The libraries were sequenced on a HiSeq2500/NextSeq500 in either paired end or single end mode. De-multiplexing and fastq file conversion were performed using blc2fastq (v.1.8.4/v2.19.1) for all libraries. After sorting, the tags and the A-overhang base were trimmed (7 bp in total). Reads from ChIP-Seq libraries were mapped using bowtie2[4] (v. 2.2.8), and filtered for uniquely mapped reads. The genome builds and annotation used for all *Drosophila* samples was BDGP6 (ensemble release 84). Peak calling was performed using macs2[5] (v 2.1.1-20160309). Further processing was done using R and Bioconductor packages. Input normalized bigwig tracks were produced using Deeptools [6] (v. 2.2.3). Heatmaps and input normalized tracks were produced using Deeptools (v. 2.2.3). Metagene profiles were produced using NGS plot[6] (v. 2.61) and Deeptools (v. 2.2.3) after input and "spike-in" normalization for libraries containing spike ins Spike in normalization was performed according to the previously published method[1].

To assign the target genes bound by $m^6A$ complex components, peaks were called using MACS2 with default cut-off and 5% FDR. The resulting peaks were annotated with the ChIPseeker package on Bioconductor, using nearest gene to the peak summit as assignment criteria. The intersection of genes bound by all $m^6A$ writer components, i.e. Mettl3-HA, Mettl14-HA, and Fl(2)d-HA, was defined as $m^6A$ complex bound.

**4sU-Seq:** The libraries were sequenced with a read length of 50 bp in paired end mode. Mapping was performed using STAR[7] (v. 2.5.1b) and ENSEMBL release 84 for Drosophila Multimapped reads were filtered out, and uniquely mapped reads to the transcript was considered.

**Calculation of Polymerase Release Ratio (PRR):** Polymerase release ratios (PRRs) were calculated as follows: for each gene, the TSS region was defined as 250 bp upstream to 250 bp downstream of the TSS. The gene body was defined as 500 bp downstream of the TSS to 500 bp upstream of the TES. The PRR ratio was calculated as the log2 ratio between the enrichment in the downstream region towards the enrichment at the TSS. For each gene, the TSS with the highest average signal around the TSS in the Control condition was selected. Enrichment calculations were based on the enrichment over the input (ChIP-Seq). The TSS reference was taken from ENSEMBL release v 84. Genes with a length smaller than the required length for the calculation were excluded.

**Machine Learning for binding prediction:** Classification samples were created by binning on a 1kb resolution. Bins are non-overlapping genomic loci that are defined as either "bound (1)" or "unbound (0)" depending on the "presence" or "absence" of a signal defined by the given m6A complex component, protein complex, or histone mark. In case of the micro-array data, when multiple regions were mapped to the same bin, the average m-value of these regions was assigned to this bin. Afterwards bins with an average m-value lower than 1.5 were assigned to unbound. Given one of the three class features Mettl3, Mettl4, or Fl(2)d all bound bins were collected and the same amount of unbound bins randomly sampled, in order to create balanced datasets. Classification features were defined by the Pol II and Ser2P datasets from this study and published datasets of GAF and M1BP together with the histone modification datasets in S2R+ cells from modENCODE consortium. To investigate whether the binding of a class defining component can be predicted by the classification features, the Random Forest algorithm with default settings but 1000 estimators was applied to create a classification model that was evaluated by the receiver operating characteristics curve (ROC) and precision recall curve (PRC) within a stratified ten-fold cross-validation.

**$m^6$A-miCLIP analysis:** Sequencing qualities were checked for all reads using FastQC (version 0.11.5) (https://www.bioinformatics.babraham.ac.uk/projects/fastqc/). Afterwards, reads were filtered based on sequencing qualities (Phred score) of the barcode region. Individual samples were processed using the CLIP Tool Kit (CTK) v1.0.9 [8]. We largely followed recommended user guide lines specific to CTK iCLIP data analysis as described here (https://zhanglab.c2b2.columbia.edu/index.php/ICLIP_data_analysis_using_CTK).
We further filtered identified CIMS and CITS to be reproducible in at least 2 out of 4 replicate $m^6$A-immunoprecipitation samples and not present in identified CITS from the input control sample for each aligner separately. Together, we considered a set of 13024 C-to-T conversions CIMS and AC truncation CITS across 2464 genes for our final S2 cell miCLIP data set. CITS site were annotated as described before [9]. For representation purposes we simplified the annotation categories. All CITS not annotated to 5'UTR, CDS, 3'UTR or intron were summarized in the category 'other'.

**Quantification and statistical analysis:** For gene expression quartile classification, all of the expressed genes in S2R+ cells in control condition were divided into equally sized quartiles according to the expression. *P*-values for significance of the associations were derived from Fisher's exact test. For PRR quartile classification, all the RNAP II binding on gene body vs promoter (PRR) in S2R+ cells were divided into four equal sized quartiles from low to high level. P-values for significance of the associations were derived from Fisher's exact test. For comparisons of the distribution of different classes we used ANOVA. t-Test were used to test the significance of tethering assays. The Kolmogorov-Smirnov test were used for testing the statistical significance for the distribution of calculated polymerase release ratio (PRR).

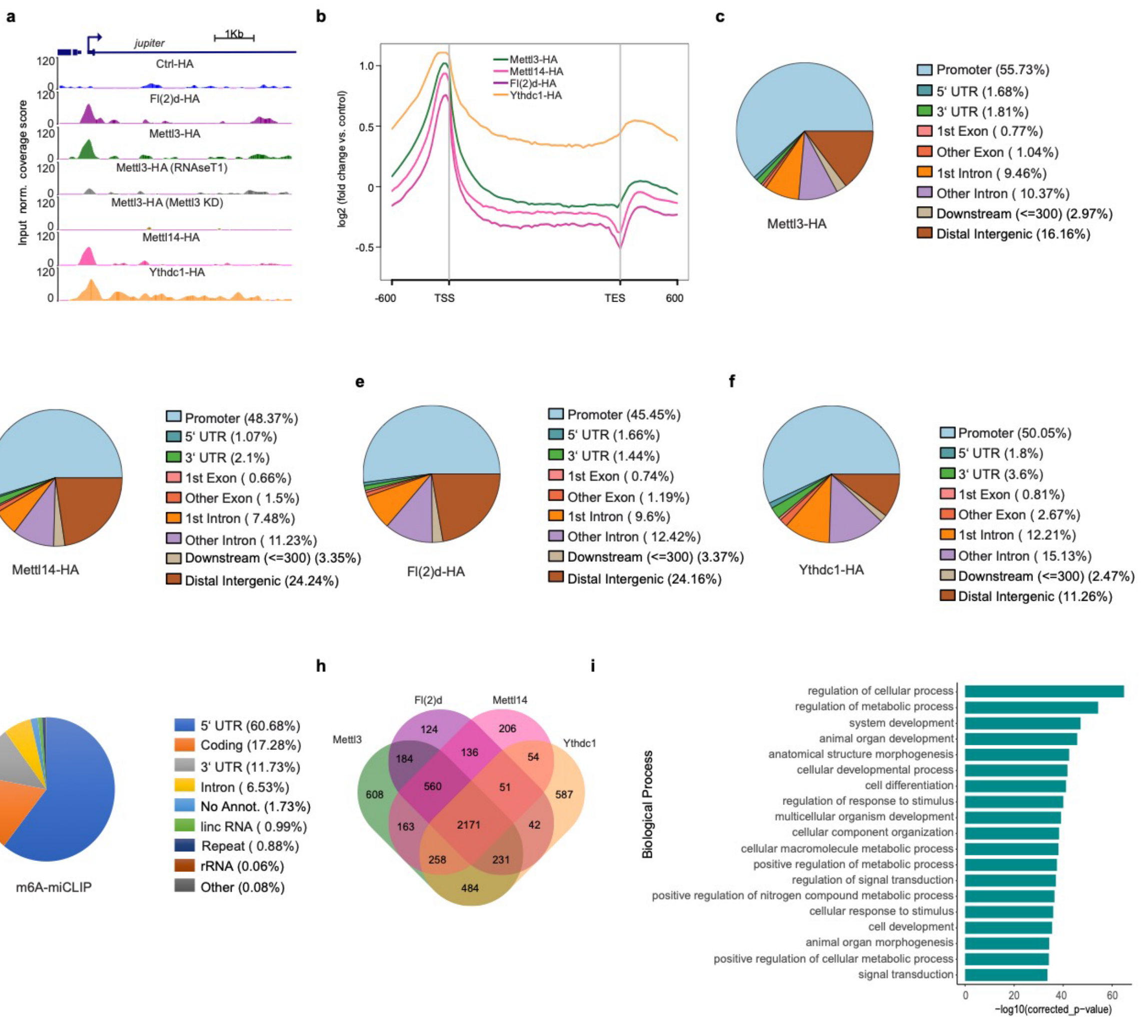


**Fig. S1. The m$^6$A complex preferentially gets recruited to promoter and is dependent on active transcription**

**(a)** Input normalized and replicate averaged track examples of ChIP-Seq experiments from S2R+ cell extracts transfected with HA-tagged m$^6$A complex components, along with the control. Shown here is recruitment of Fl(2)d, Mettl3, Mettl14 and Ythdc1 to *Jupiter*. **(b)** Averaged metagene profiles with standard error of mean from two independent biological replicates of ChIP-Seq performed with HA-tagged m$^6$A complex components. Log2 fold changes against input control are shown on Y-axis while X-axis depicts scaled genomic coordinates. **(c-f)** Pie charts depicting distribution of genomic features of annotated peaks of the components of m$^6$A complex. Shown are the pie charts for Mettl3 **(c)**, Mettl14 **(d)**, Fl(2)d **(e)** and Ythdc1 **(f)**. Note that the majority of the peaks are at the promoters of the genes for individual components of m$^6$A complex. **(g)** Pie chart depicting distribution of genomic features of m$^6$A miCLIP from S2R+ cells. The majority of the peaks are at the 5' UTR**.** **(h)** Venn diagram showing the overlap between genes bound by m$^6$A complex: writers and nuclear reader. **(i)** Gene ontology analysis showing enriched biological processes among the set of genes co-bound by m$^6$A writer complex.

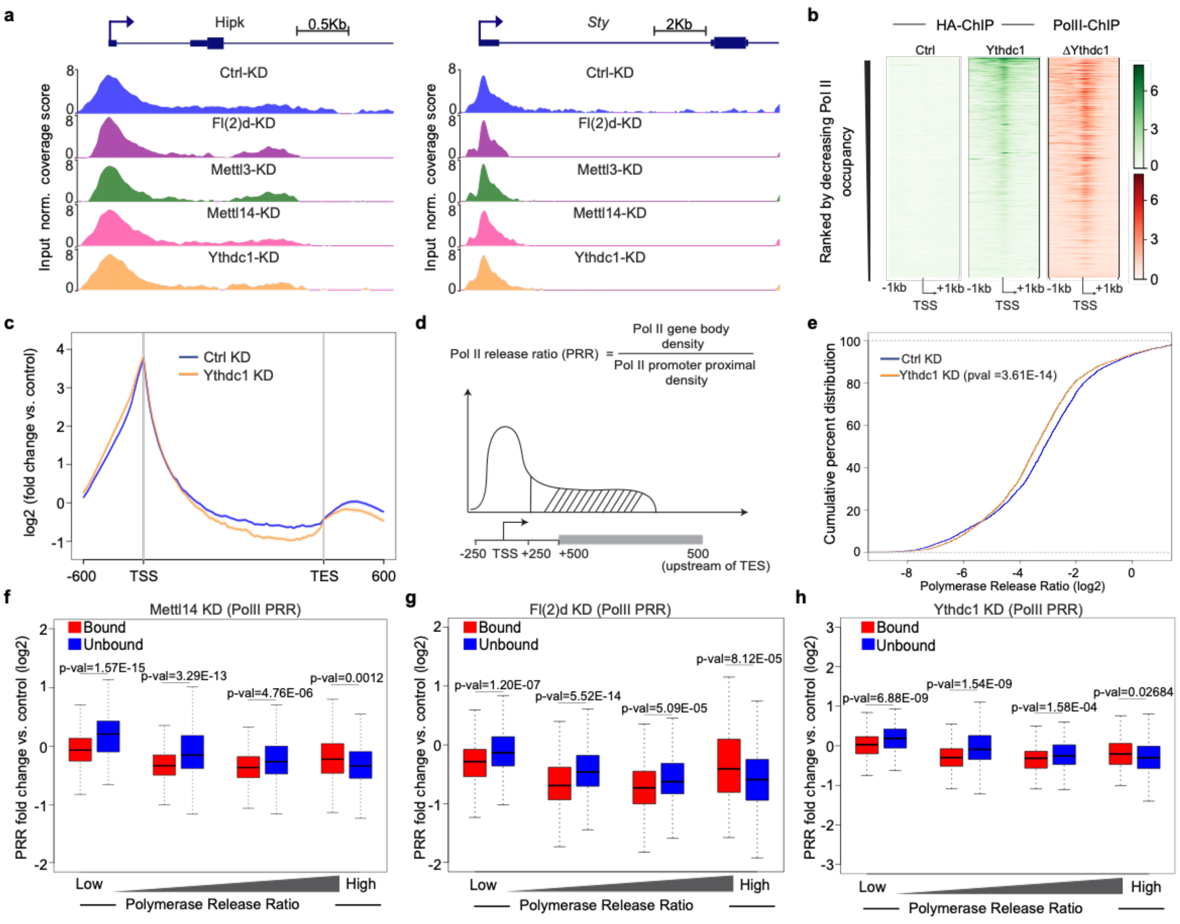


**Fig. S2. The m$^6$A complex together modulates promoter proximal pause release of Pol II**

**(a)** Track examples of total Pol II ChIP-Seq from S2R+ cell extracts, after either control or m$^6$A complex knockdowns, as indicated in the labels. The tracks are average of two independent biological replicates after input and "“spike-in”" normalization. **(b)** Heatmaps of HA-tagged Ythdc1 and change in total Pol II relative to control condition, upon Ythdc1 knockdown, centered at the TSS (-1kb to +1kb). Rows indicate all the genes bound by Pol II and are sorted by decreasing Pol II occupancy. The color labels to the right indicate the levels of enrichment. **(c)** Metagene profiles of averaged total Pol II occupancies from two independent biological replicates after "“spike-in”" normalization in control and Ythdc1 depleted cells along with standard error of mean for all the Pol II bound genes. Log2 values against input control is on Y-axis. **(d)** Schematic representation of Pol II Release Ratio calculation. **(e)** The empirical cumulative distribution function (ECDF) plot of computed PRR in control and Ythdc1 knockdown conditions, after “spike-in” normalization. P-value is derived from two-sample Kolmogorov-Smirnov test. **(f-h)** Boxplot showing change in PRR upon knockdowns across different PRR quartiles, further classified based on binding by $m^6A$ writer complex. For PRR quartile classification, all the expressed genes in S2R+ cells were divided into four equal sized quartiles according to the level of PRR, from low to high level. Shown here are the change in PRR upon Mettl14 **(f)**, Fl(2)d **(g)**, and Ythdc1 **(h)** knockdowns. P-values for significance of the associations, derived from Fisher’s exact test, are shown on top of the histograms.

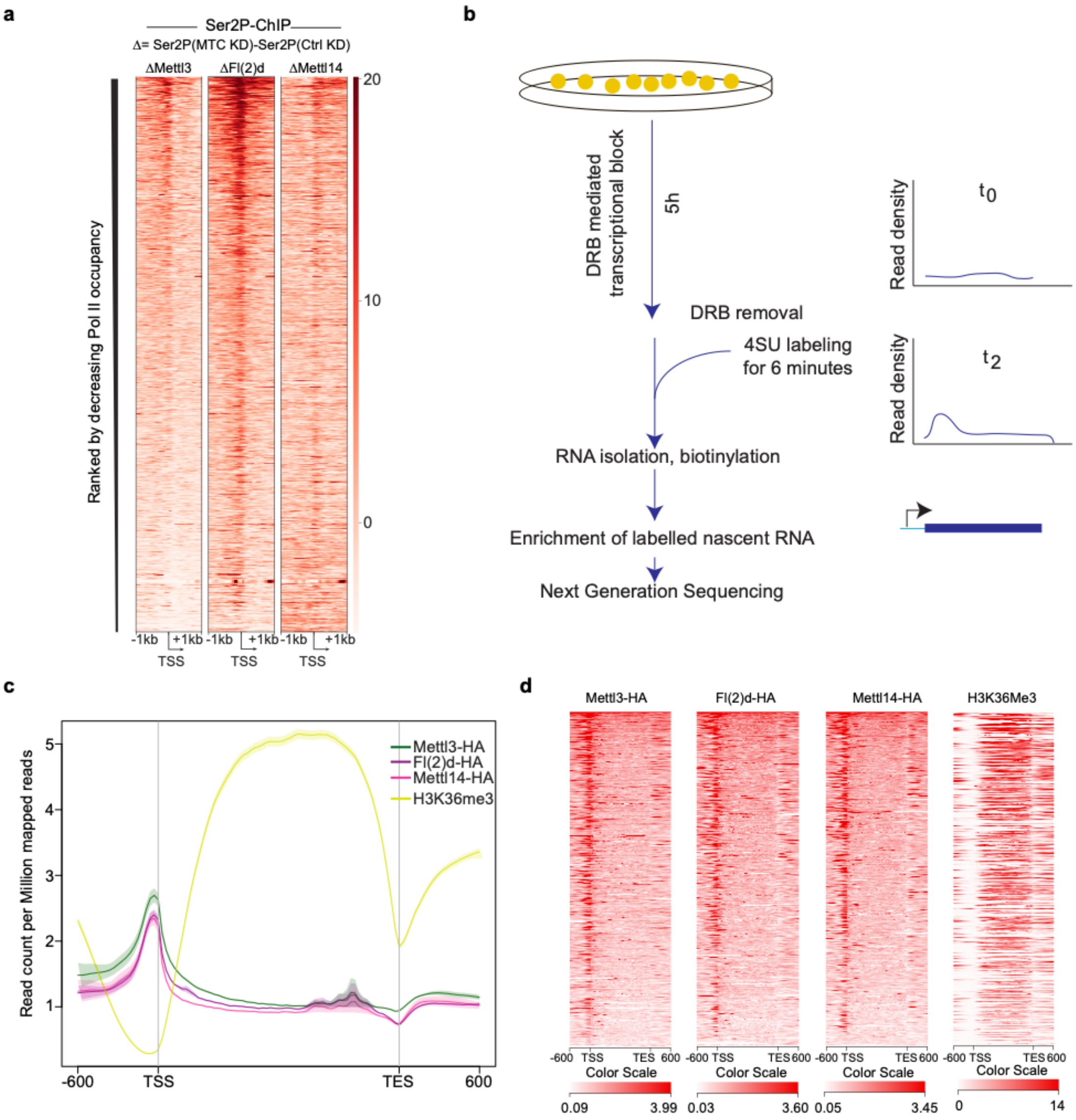


**Fig. S3. The effect of m$^6$A MTC on transcriptional elongation is not mediated through H3K36me3**

**(a)** Heatmaps of change in Ser2P occupancy upon indicated knockdowns in S2R+ cells, centered at the TSS (-1kb to +1kb). Rows indicate all the genes bound by Pol II and are sorted by decreasing Pol II occupancy. The color labels to the right indicate the levels of enrichment. **(b)** Schematic representation of DRB-4SU-Seq approach. (**c**) Metagene profiles of ChIP-seq performed with HA-tagged m$^6$A complex components and with H3K36me3 in S2R+ cells, with standard error of mean. Read counts per million of mapped reads is shown on Y-axis while X-axis depicts scaled genomic coordinates. **(d)** Heatmaps of HA-tagged m$^6$A complex components and H3K36me3 over the genebody. Rows indicate all the genes bound by Pol II and are sorted by decreasing Pol II occupancy. The color labels to the right indicate the levels of enrichment.

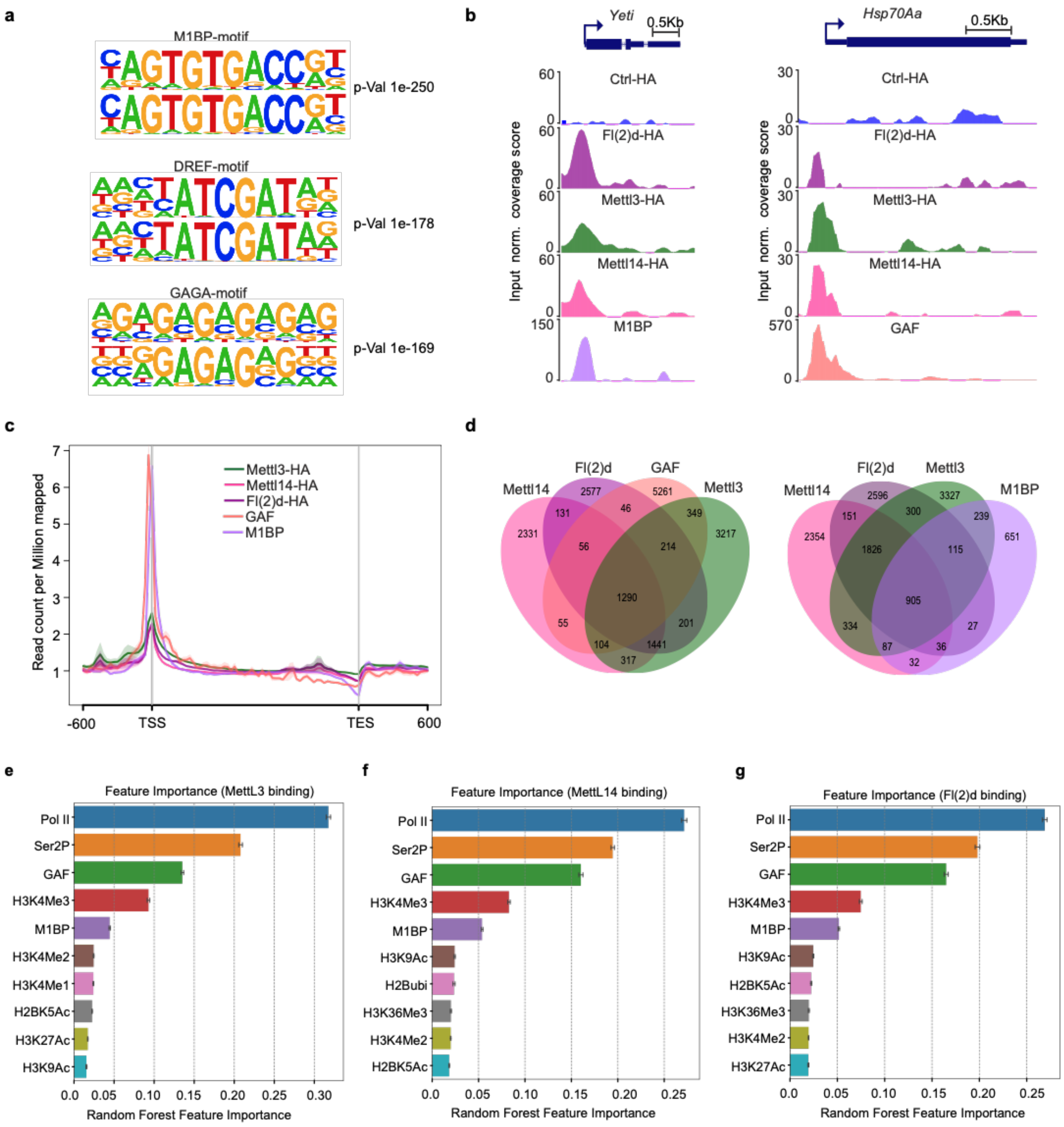


**Fig. S4. The GAF, DREF, and M1BP motifs are overrepresented in the m$^6$A writer complex bound peaks**

**(a)** Motif logo of the factors whose consensus sequences were significantly overrepresented in the peaks shared by all the components of m$^6$A writer complex. The p-values for the enrichment of the motif is displayed next to the motif logo. The motif analysis was performed with the Homer package, see methods section for further details. **(b)** Input normalized and replicate averaged track examples of ChIP-Seq experiments from S2R+ cell extracts transfected with HA-tagged m6A complex components, along with the control. Shown here is recruitment of Fl(2)d, Mettl3, and Mettl14 along with the co-binding of M1BP and GAF, on the indicated genes. The publicly available GAF and M1BP data were normalized to their respective IgG control. **(c)** Metagene profiles with standard error of mean ChIP-seq performed with HA-tagged m$^6$A complex components and with the publicly available ChIP-seq datasets of GAF and M1BP in S2R+ cells. Read counts per million of mapped reads is shown on Y-axis while X-axis depicts scaled genomic coordinates. **(d)** Venn diagram showing the overlap between genes bound by m$^6$A writer complex and GAF, and with M1BP. **(e-g)** Histograms showing the Random Forest feature importance for different datasets used for predicting the recruitment of Mettl3 **(e)**, Mettl14 **(f)**, and Fl(2)d **(g)**. Shown are the ten most important features, error bars indicate the standard deviation from the mean across the ten-fold cross-validation.

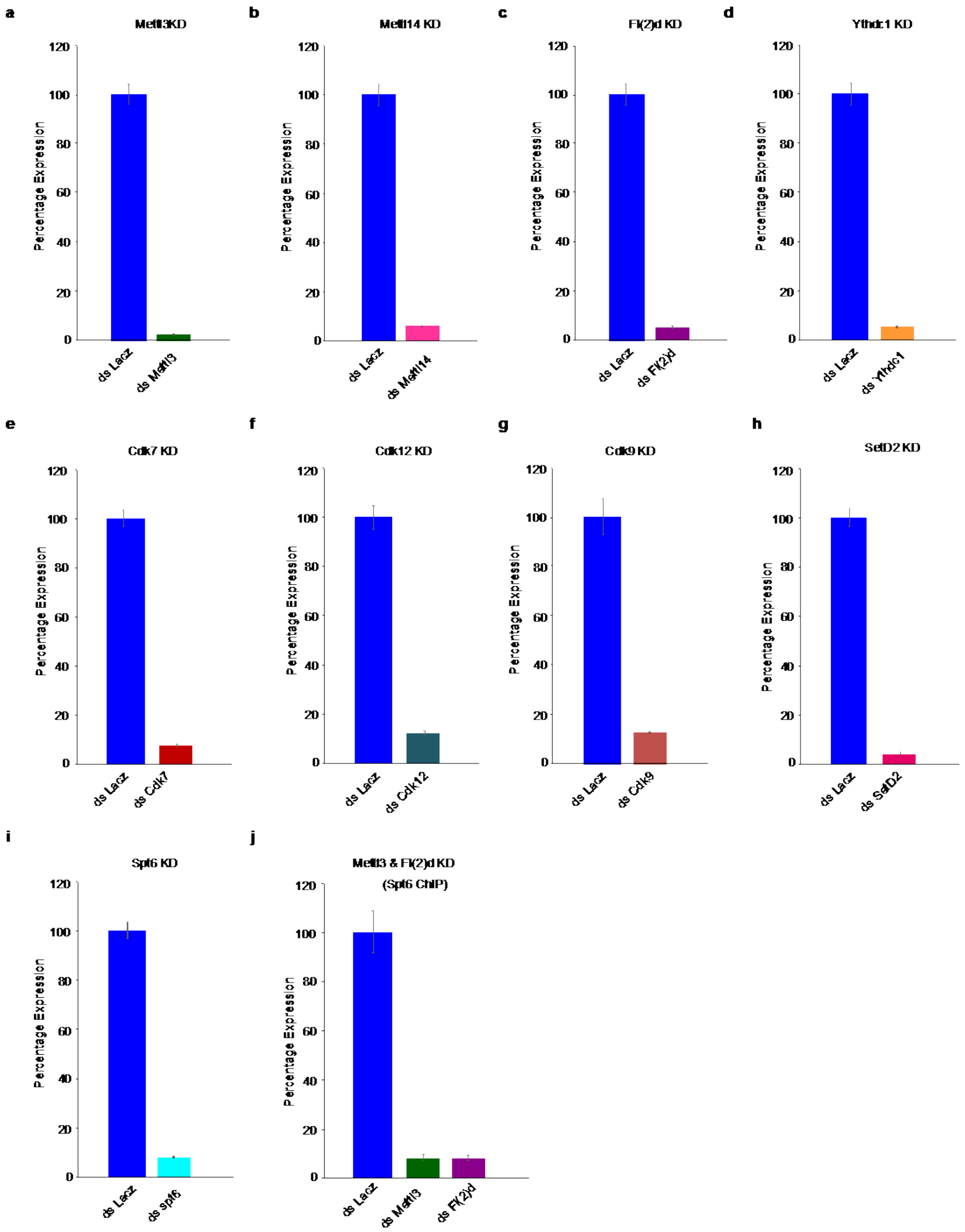


**Fig. S5. The RT-qPCR analysis of the knockdown efficiencies**

RT-qPCR analysis showing the efficiency of knockdown for various conditions. Shown here are knockdown efficiencies for Mettl3 **(a)**, Mettl14 **(b)**, Fl(2)d **(c)**, Ythdc1 **(d)**, Cdk7 **(e)**, Cdk12 **(f)**, Cdk9 **(g)**, SetD2 **(h)**, Spt6 **(i)**, and double knockdown of Mettl3 and Fl(2)d **(j)**.

**Table S1. Table depicting potential m$^6$A RNA nuclear interactors**

| Prot. FDR conf. | Accession | Gene Name | Function | Exp. q-value | PEP Score | Coverage | Peptides | PSMs | Uniq. Pept. | Pep Seq HT |
|---|---|---|---|---|---|---|---|---|---|---|
| High | P25823 | tud | transport/mRNA localiz. | 0 | 34,9 | 4,7 | 7 | 9 | 1 | 7 |
| High | Q8IMV6 | Saf-B | mRNA binding/splicing | 0 | 13,2 | 6,5 | 4 | 7 | 4 | 4 |
| High | Q9VUH8 | Tdrd3 | RNA binding | 0 | 12,1 | 9,0 | 5 | 5 | 5 | 5 |
| High | Q7JQT9 | CG1671 | RNA binding | 0 | 11,9 | 6,2 | 3 | 3 | 3 | 3 |
| High | Q9W3L1 | Upf2 | RNA binding/NMD | 0 | 7,0 | 1,7 | 2 | 2 | 2 | 2 |
| High | Q9VV73 | nxf2 | nuclear mRNA export | 0 | 6,9 | 3,6 | 2 | 2 | 2 | 2 |
| High | Q9W410 | Pf1 | chromatin binding | 0 | 6,0 | 3,5 | 2 | 2 | 2 | 2 |
| High | P21525-3 | kay isoB | TF | 0 | 24,5 | 14,1 | 5 | 17 | 1 | 5 |
| High | A0A0B4LHU | kay isoG | TF | 0 | 23,0 | 10,2 | 5 | 17 | 1 | 5 |
| High | Q9VHG5 | Ibf1 | Chromatin insulator | 0 | 11,3 | 24,0 | 3 | 3 | 3 | 3 |
| High | Q8INP9 | CG11980 | | 0 | 10,3 | 9,7 | 2 | 2 | 2 | 2 |
| High | O02193 | mof | histone acetyl transferase | 0 | 7,0 | 4,1 | 2 | 2 | 2 | 2 |
| High | Q9W3X8 | CG3198 | mRNA binding/splicing | 8,9526E-05 | 4,7 | 4,8 | 2 | 2 | 2 | 2 |
| High | Q7JVK6 | trsn | RNA binding/RNA interference | 8,9526E-05 | 4,1 | 13,2 | 2 | 2 | 2 | 2 |
| High | N0D8I3 | Spt6 | chromatin binding/PolII binding | 8,9526E-05 | 4,0 | 2,3 | 3 | 3 | 3 | 3 |
| High | Q8SWT2 | Rhau | RNA helicase | 0,00010939 | 4,8 | 2,5 | 2 | 3 | 1 | 2 |
| High | Q9VYY7 | HP5 | heterochromatin | 0,00017283 | 3,0 | 5,4 | 2 | 2 | 2 | 2 |
| High | Q8INB9 | Akt1 | Protein kinase | 0,00017283 | 2,9 | 6,5 | 2 | 3 | 2 | 2 |

**Table S2. Primers used for cloning**

| Name | Sequence |
|---|---|
| Box B Fwd KpnI | NNNGGTACCACTACCAAACTGGGGATTCCTG |
| Box B Rev KpnI | NNNGGTACCGACCTCGAGATAATATCCTCGATAG |
| Mettl3 Fwd | ATCATGGCAGATGCGTGGGACATAAAATC |
| Mettl3 Rev | NNNNGCGGCCGCCTACTTTTGTATTCCATTGATCGACG |
| eGFP EcoRV Fwd | NNNNGATATCATGGTGAGCAAGGGCGAGGA |
| eGFP NotI Rev | NNNNGCGGCCGCTTACTTGTACAGCTCGTCCATGCC |
| MettL3 EcoRV Half F | ATCATGGCAGATGCGTGGGACATAAAATC |
| MettL3 EcoRV mut R | ATCCCCGGCAGCACCGGCCATCACCACAGCGAAC |

**Table S3. Primers used for knockdown experiments**

| Name | Sequence |
|---|---|
| LacZ-T7 Fwd | TAATACGACTCACTATAGGGCAGGCTTTCTTTCACAGATG |
| LacZ-T7 Rev | TAATACGACTCACTATAGGGCTGATGTTGAACTGGAAGTC |
| Cdk9-T7 Fwd | TAATACGACTCACTATAGGGCTACCAAACCCGGTCAATCATAC |
| Cdk9-T7 Rev | TAATACGACTCACTATAGGGATCCTGCACCGAGACATGAAAG |
| Cdk7-T7 Fwd | TAATACGACTCACTATAGGGACAAGGCGCGACACGGTGA |
| Cdk7-T7 Rev | TAATACGACTCACTATAGGGCTTTCAATCACATCTTTAATAGATCATCC |
| Mettl3 T7 F | TTAATACGACTCACTATAGGGAGACAGCCTGGAGATGGTGAACT |
| Mettl3 T7 R | TTAATACGACTCACTATAGGGAGATCAGGCACTCGACTTTTGTG |
| Mettl3 T7 F2 | TTAATACGACTCACTATAGGGAGAGCACTGGTCATTGGCTAAAC |
| Mettl3 T7 R2 | TTAATACGACTCACTATAGGGAGACATCTGGATAACGCTTCTGGA |
| Mettl14 T7 F | TTAATACGACTCACTATAGGGAGATAATCAGAATGCCGCCACTA |
| Mettl14 T7 R | TTAATACGACTCACTATAGGGAGATGATCAGTTCCCTCAACTTGG |
| Mettl14 T7 F2 | TTAATACGACTCACTATAGGGAGATAACAGTAGTCGCCAGCTGAATTCC |
| Mettl14 T7 R2 | TTAATACGACTCACTATAGGGAGACTCAGCAAAGTAGGTCTGGTATAGC |
| fl(2)d F T7 | TTAATACGACTCACTATAGGGAGACGGACTCTAATACCGGCAAA |
| fl(2)d R T7 | TTAATACGACTCACTATAGGGAGAGGTGTTGTCACTGGCTGATG |

| | |
|---|---|
| fl(2)d T7 F2 | TTAATACGACTCACTATAGGGAGAGCCCCTATGAAAAGCAACAG |
| fl(2)d T7 R2 | TTAATACGACTCACTATAGGGAGAGGATCGAGCAGAGCAGTG |
| Ythdc1 T7 F | TTAATACGACTCACTATAGGGAGACGAATCGAATGGTGGAGACT |
| Ythdc1 T7 R | TTAATACGACTCACTATAGGGAGACCGTGTGTCTCGGAATAGGT |
| Ythdc1 T7 F2 | TTAATACGACTCACTATAGGGAGAACGGAGCACCACACCATC |
| Ythdc1 T7 R2 | TTAATACGACTCACTATAGGGAGAGTCCCGATTCCGATAGCC |
| M1BP T7 Fwd | TAATACGACTCACTATAGGGCTGGAGAACTACGGTTGCCT |
| M1BP T7 Rev | TAATACGACTCACTATAGGGTCGCAGCCAAATTGCTTGT |
| GAGA T7 Fwd | TAATACGACTCACTATAGGGAATACACCATGCAAGCATCC |
| GAGA T7 Rev | TAATACGACTCACTATAGGGCGCCAAGCACCTGCTGCA |
| DREF T7 Fwd | TAATACGACTCACTATAGGGACCAGCGTCGCCCGTGGC |
| DREF T7 Rev | TAATACGACTCACTATAGGGGATCAAAGAGTCGATCTTGTGTG |
| Spt6 T7 Fwd | TAATACGACTCACTATAGGGATGAGTACGATGAGGATCTGGG |
| Spt6 T7 Rev | TAATACGACTCACTATAGGGATACCGCATAGGGACTGCTG |

**Table S4. Primers used for quantitative PCR (qPCR) experiments**

| Name | Sequence |
|---|---|
| M1BP_qPCR_Fwd | CCGCAAGTACCACAGACGTA |
| M1BP_qPCR_Rev | ACAACTTCCGAGCTGAAACG |
| GAGA_qPCR_Fwd | CACTTGCTGCACCTGTTGTT |
| GAGA_qPCR_Rev | GGTTCAATCCCTGCACACTT |
| Box B 5' F | GCCCCTCGACTAGTCCAAAT |
| Box B 5' R | CCGGGCCTTTCTTTATGTTT |
| Act 5' | ACACAA AGCCGCTCCA TCAG |
| Act5' F | CATTGCGGCTGATAAGGTTT |
| Act5' R | GCTTTGTGTCGGGAGGAGTA |
| Luc 5' F | TACCAACATGGCCGAAGAC |
| Luc 5'R | GGAACCAGGGCGTATCTCTT |
| Luc 5' F 2 | TTGTTTCCAAAAAGGGGTTG |
| Luc 5'R2 | ACGTGTACATCGACTGAAATCC |
| Luc GB F | TGGGTTACCTAAGGGTGTGG |
| Luc GB R | AAAACCGTGATGGAATGGAA |
| Luc GB F 2 | TCGCCAAAAGCACTCTGATT |
| Luc GB R 2 | CTGGAAGATGGAAGCGTTTT |
| Luc 3' F | GTAACAACCGCGAAAAAGTTG |
| Luc 3' R | GCCCTTCTTGGCCTTTATG |
| Luc 3' F2 | GGGACGAAGACGAACACTTCT |
| Luc 3' R2 | CGAAGATGTTGGGGTGTTG |

| | |
|---|---|
| Ren 5' F | AGTGGTGGGCCAGATGTAAA |
| Ren 5'R | AATAAGAAGAGGCCGCGTTA |
| Ren GB F | TTATTGAATCGGACCCAGGA |
| Ren GB R | CATCAGGTGCATCTTCTTGC |
| Ren 3' F | TTATTGAATCGGACCCAGGA |
| Ren 3' R | CATCAGGTGCATCTTCTTGC |
| Cdk12-F | TTCGGGAACTGGTACCTCTG |
| Cdk12-R | CGGCGATGATGATAGTGATG |
| Cdk7-F | TGTGGAAAAGATCCGTGTCA |
| Cdk7-R | GTCACACGCTGGTATCGTTC |
| Primer- Neg Fwd (Pol II ChIP) | CCATTAATCGAGGGCTGAAA |
| Primer- Neg Rev (Pol II ChIP) | TTGGGGCATAAACAGAGGAC |
| Cdk9 qPCR Fwd | TCGTGTTCGCAGAAATCAAA |
| Cdk9 qPCR rev | TAAAGCACGAGAACGTGGTG |
| PolII Neg-Fwd | AGCAATGTCGCTTCACACAC |
| PolII Neg-Rev | CTTGCGCCTAAGCTATTTGG |
| Mettl13 Fwd | AAGGAACTCGTTGAGGCTGA |
| Mettl13 Rev | CACCTGTGTGGAGACAATGG |
| Mettl14 Fwd | AAGCGTCGTTTGCTTTTAGC |
| Mettl14 Rev | GCATTACCCAAAGCCTTTTTC |
| Ythdc1 Fwd | GGCTCGAGTTATGCGAGAAA |
| Ythdc1 Rev | GGTGGTCGTGATTTGATCCT |
| Rpl15 Fwd | AGGATGCACTTATGGCAAGC |

| | |
|---|---|
| Rpl15 Rev | GCGCAATCCAATACGAGTTC |
